# Glycosaminoglycans and glycoproteins influence the elastic response of synovial fluid nanofilms on model oxide surfaces

**DOI:** 10.1101/2021.11.19.469284

**Authors:** Amar S. Mann, Ariell M. Smith, Joyce O. Saltzherr, Arvind Gopinath, Roberto C. Andresen Eguiluz

## Abstract

Synovial fluid (SF) is the natural lubricant found in articulated joints, providing unique cartilage surface protecting films under confinement and relative motion. While it is known that the synergistic interactions of the macromolecular constituents provide its unique load-bearing and tribological performance, it is not fully understood how two of the main constituents, glycosaminoglycans (GAGs) and glycoproteins, regulate the formation and mechanics of robust load-bearing films. Here, we present evidence that the load-bearing capabilities, rather than the tribological performance, of the formed SF films depend strongly on its components’ integrity. For this purpose, we used a combination of enzymatic treatments, quartz crystal microbalance with dissipation (QCM-D) and the surface forces apparatus (SFA) to characterize the formation and load-bearing capabilities of SF films on model oxide (i.e., silicates) surfaces. We find that, upon cleavage of proteins, the elasticity of the films is reduced and that cleaving GAGs results in irreversible (plastic) molecular re-arrangements of the film constituents when subjected to confinement. Understanding thin film mechanics of SF can provide insight into the progression of diseases, such as arthritis, but may also be applicable to the development of new implant surface treatments or new biomimetic lubricants.

## INTRODUCTION

Despite intense research efforts to elucidate the molecular mechanisms that allow articular joints to support high stresses and deformations over a person’s life span, it remains unclear what role synovial fluid (SF) plays in load-bearing and boundary lubricating conditions. SF is a complex fluid composed of a mixture of biomolecules and ions, such as glycosaminoglycans (*e.g*., hyaluronan), glycoproteins (*e.g*., lubricin), seric proteins (*e.g*., albumin), lipids (*e.g*., DPPC), among others. These give rise to a highly viscous liquid in homeostasis, with various identified biochemical[1] and biomechanical[2] functions. SF components adsorb to the articular cartilage or implant surfaces and form films that arrest cell attachment[3,4] and maintain surfaces separated at high contact pressures. This prevents adhesion,[5] and when the surfaces are in relative motion, provides wear protection and lubrication.[6,7]

The ability of specific SF components to adsorb to surfaces is crucial to build and control the supramolecular assembly of an effective load-bearing, wear-protecting, and lubricating film. Without it, molecules would be squeezed out from the junction at the high contact (normal) loads experienced during locomotion (~10-15 MPa).[8] For example, hyaluronan, a major component of SF, has been shown to provide remarkable wear protection only when strongly bound (chemisorbed) to model mica substrates, expelled and poorly protecting against wear when weakly bound (physisorbed).[9–11] Molecular complexation has been demonstrated to control the mechanical response of SF components. For instance, single-molecule stretching studies using magnetic tweezers force spectroscopy exemplified that the degree of glycosylation regulates tension in the hyaluronan-aggrecan bottlebrush complex,[12] abundant in SF, and is suggested to impact cartilage normal load bearing at the mesoscale.[13] Additionally, concentrations of SF components can tune the conformations adopted by biomolecules due to crowding effects. That is the case for isolated lubricin, which adopts different conformations when adsorbed to mica surfaces depending on its concentration.[14] Combined, it is clear that each of the above mentioned parameters (*i.e*., adsorption to surfaces, complexation, conformation) are crucial for load-bearing films’ proper function. A biochemical imbalance will invariably result in a biomechanical imbalance as well. This is evidenced in rheology studies of synovial fluids obtained from patients with various autoimmune or trauma related joint pathophysiology, in which downregulation of SF component concentrations or shift in molecular weight distributions lead to altered viscosity and viscoelastic responses.[4,15]

Many SF studies have focused on the lubricity and wear protection properties of isolated components, such as lubricin,[14,16] hyaluronan,[10,11] or lipids.[17,18] Higher-order mixtures studies, such as hyaluronan and lubricin,[19–21] hyaluronan and phospholipids,[22–25] lubricin and galectins,[26] or hyaluronan and aggrecans,[27,28] combined with a top-down approach in which different components have been enzymatically digested,[29,30] emphasize that it is the synergistic interactions that provide SF’s unique load-bearing, wear protecting, and lubricating properties. Still, less is known about the formation of SF films on surfaces and film properties, such as viscoelasticity and load-bearing capabilities. The present study provides evidence that the load-bearing capabilities of the formed SF films depends strongly on the integrity of its components. Particularly, we focus on glycosaminoglycans (GAGs) and glycoproteins’ roles in the formation kinetics, viscoelasticity, and load-bearing mechanics of protecting SF films. For this purpose, we used a combination of quartz crystal microbalance with dissipation (QCM-D) and the surface forces apparatus (SFA) to characterize the formation and load-bearing properties of SF films on model oxide (silica and mica) surfaces and elucidate the SF film’s structural changes under different enzymatic treatments and dilutions. We find that the absence of either GAGs or proteins dysregultes the load-bearing capabilities. Therefore, the findings reported can be of relevance to understand the formation of protecting films on cartilage or implant surfaces. Furthermore, we emphasize the importance of load-bearing properties of SFs under confinement in addition to the tribological characterization, to fully understand the supramolecular assemblies and synergistic interactions of SF components.

## EXPERIMENTAL SECTION

### Synovial fluids

All experiments performed with synovial fluid (SF) were in agreement with the procedures and guidelines provided by the biosafety committee at the University of California Merced.

Pooled bovine SF (Lampire Biological Laboratories, 8600853) was used to prepare working aliquots, kept frozen (*T* = −80 °C) and thawed the day of the experiment. SF was first centrifuged for 5 min at 6000 rpm for all experiments to remove cell debris and large tissue aggregates, and suspension moved to a new tube. This condition is referred to from now on as nontreated SF. To cleave GAGs in SF, centrifuged and undiluted SF was treated with 2 μL of 1.25 mg/mL hyaluronidase (HAase) from bovine testes (Sigma, H3506) for every 50 μL of SF, following established protocols. [7,30] SF plus the hyaluronidase were placed on a shaker for 1.5 to 2 hrs., followed by a second centrifugation step, 5 min at 6000 rpm, and the suspension transferred to a new tube. This condition is referred to from now on as SF+HAase. To digest proteins present in SF, centrifuged and undiluted SF was treated with 2 mg/mL trypsin from bovine pancreas, tosyl phenylalanyl chloromethyl ketone (TPCK) treated and salt-free (Sigma, T1426) stock solution, from which 25 μL were added for every 1 mL SF, following established protocols. [7,30] SF plus trypsin were placed in an incubator at 37 °C for at least 2 hrs. There was no need for a second centrifugation step for trypsin-treated SF, as no supernatants were ever found. This condition is referred to from now on as SF+Trypsin.

Nontreated SF, SF+HAase, and SF+Trypsin were diluted to the following percentages using phosphate buffered saline (PBS) (Gibco, 10-010-023): 1%, 5 %, 50%, and 100% (undiluted) solutions for QCM-D, and 5% and 100% for SFA experiments.

### Quartz Crystal Microbalance with Dissipation (QCM-D)

In QCM-D measurements, a quartz crystal oscillator is set to oscillate at its resonance frequency. The shift in frequency due to the formation of an SF adsorption layer is typically 10-200 Hz. The frequency and dissipation changes can be related to the mass oscillating with the crystal and the viscoelastic properties of the layer through various models.^35^

### Surfaces Forces Apparatus (SFA)

The SFA allows to measure normal forces (*e.g*., forces due to the surface and liquid structure, polymer, steric) between macroscopic surfaces with high force sensitivity while simultaneously monitoring the absolute surface distance (film thickness) and contact shape.

A full detailed experimental protocol with all relevant equations and theoretical modeling of QCM-D and SFA can be found in the Supporting Information.

## RESULTS

### Film formation kinetics

The changes in surface mass density (Δ*Γ*) and dissipation (Δ*d*) during the adsorption, rinsing, and post-rinsing of nontreated SF, SF+HAase and SF+Trypsin to a silica oscillator at the four tested concentrations are shown in Figure 1 and SI Figure 1. We observed, as expected, that the time to reach the maximum surface mass density has a strong dependency on the SF dilution and does weakly depend on treatment of SF. We used a kinetic model, SI eq. 3, to quantitatively obtain saturation times (*τ*) at any given concentration, summarized in SI Table 1 and SI Figure 2. For all treatments, *τ* decreased with increasing concentration. For nontreated SF, SF+HAase, and SF+Trypsin at one percent (1%), *τ* values were 3.4 ± 1.9 min, 4.0 ± 1.9 min, and 1.6 ± 0.9 min, respectively. *τ* values decreased to 2.0 ± 1.1min for nontreated SF, 2.1 ± 0.9 min for SF+HAase, 0.7 ± 0.1 min for SF+Trypsin at five percent (5%). At 50%, *τ* reached a minimum value of 0.9 ± 0.3 min, 0.9 ± 0.2 min, and 0.9 ± 0.5 min and remained practically unchanged at 100%, being 0.7 ± 0.3 min, 0.9 ± 0.3 min, 1.0 ± 0.2 min for nontreated SF, SF+HAase, and SF+Trypsin, respectively. The dissipation behavior of the films as a function of time followed an identical qualitative trend as what described for the change in surface mass density for all concentrations and treatments. Nontreated SF at 100% stands out from the rest of the conditions. Δ*Γ* and Δ*d* values decrease after reaching an almost instantaneous maxima, as shown in Figure 1(d). After the 30 minutes of experimental time, the film continues to experience molecular re-arrangements and has not reached a steady state/equilibrium. Most likely components with higher molecular weight (*MW*) and complexes are slowly pushed upward due to the oscillating surface in a phenomena that bears similarity to the vibration-induced granular segregation (or the Brazil nut effect),[37] combined with competing events of molecules with higher affinities to silica surface. Furthermore, SF+Trypsin presented an initial sharp peak, decreasing in amplitude with increasing concentration, which we attribute to fluid pressure gradients acting on the sensor, due to the lack of glycoproteins dissipating shear stresses.

**Figure 1.**
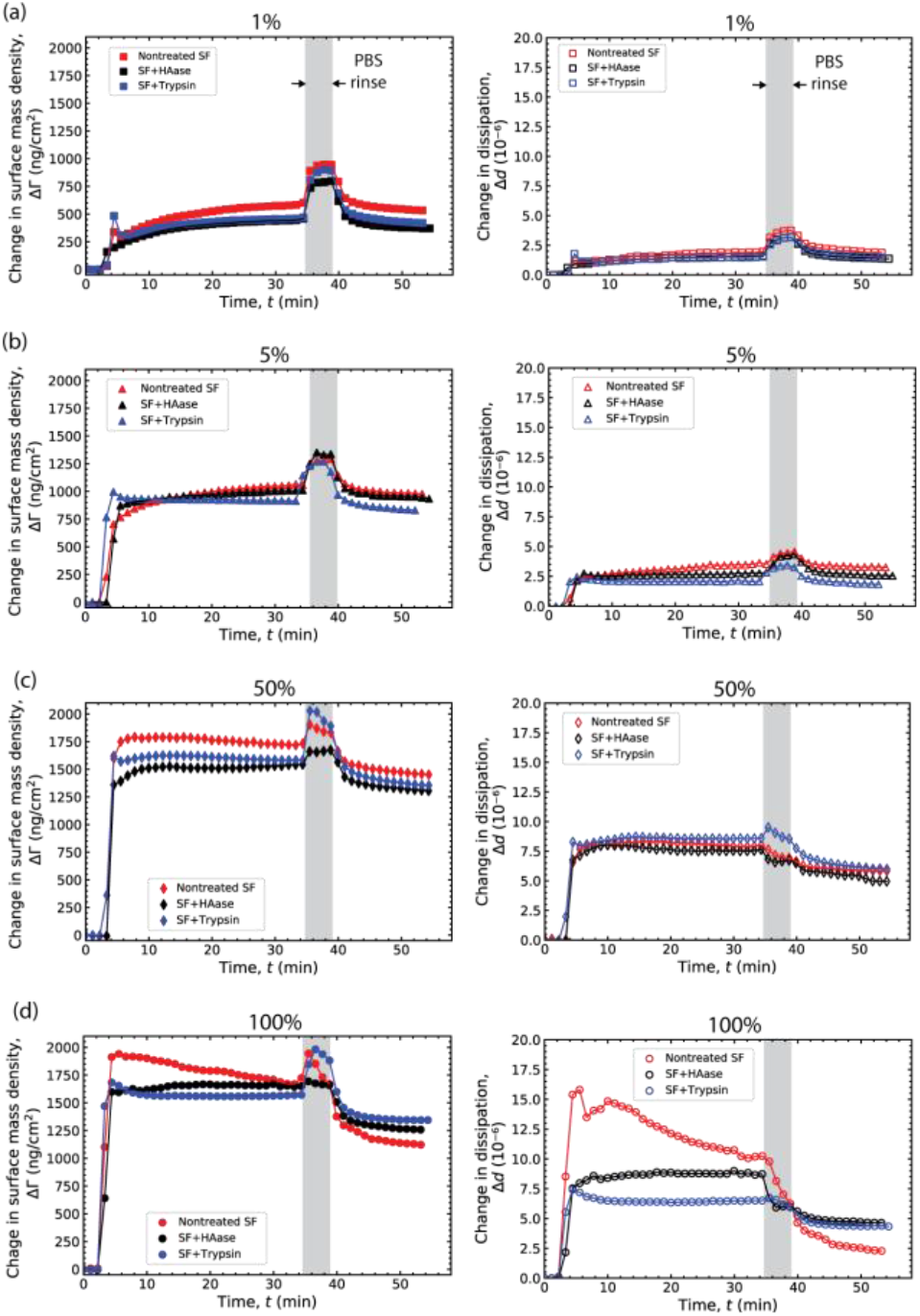
Surface mass density and dissipation change during the adsorption of SF to silica. (a) One percent (1%) in PBS, (b) five percent (5%) in PBS, (c) 50 percent (50%) in PBS, and (d) nondiluted (100%) nontreated SF (red), SF+HAase (black), and SF+Trypsin (blue). For clarity, only every 100^th^ data point is shown.

Next, we rinsed nontreated SF, SF+HAase, and SF+Trypsing films with PBS to quantify the amount of SF components strongly physisorbed to the silica oscillator surface and the change in dissipation for each condition (treatment and concentration). After rinsing with PBS, the change in surface mass density and change in dissipation did not significantly shift for the lower concentrations, one percent and five percent of any of the treatments, summarized in SI Table 1 and denoted ΔΔ*Γ*. However, rinsing with PBS did considerably shift the change in surface mass density and change in dissipation to lower values for the two higher concentrations, 50% and 100% of nontreated SF, SF+HAase, and SF+Trypsin, as further described in the following section, *Langmuir adsorption isotherms*.

These results indicate that at 50% concentration, independent of the enzymatic treatment, a full film, strongly adhered to the silica oscillator surface has formed, and that a second weakly bound layer has started to build, which was easily removed by rinsing with PBS. Again, nontreated SF at 100% presented a unique behavior, as the dissipation after the PBS rinse dramatically shifted to very low values.

### Langmuir adsorption isotherms

To describe the coverage of molecules adsorbed onto the silica oscillator and the dissipation of the films formed as a function of SF concentrations, we used the Langmuir isotherm model, SI eq. 4, for nontreated SF, SF+HAase, and SF+Trypsin before and after a PBS rinse, Figure 2. Here, we report the surface mass densities (*Γ*) and dissipation (d) rather than Δ*Γ* and Δ*d* as we report the average over the last 20 data points, illustrated in SI Figure 3. The *Γ* were similar between treatments at all tested concentrations (1%, 5%, 50%, and 100%). These findings are not surprising, as the enzymes randomly cleave b-N-acetylhexosamine-glycosidic bonds in GAGs (HAase)[38] or proteins at the carboxyl side of Lys and Arg residues (Trypsin) and we did not extract the fragments. Interestingly, we found that the maximum surface mass density was reached with at least 50% nontreated SF, SF+HAase, or SF+Trypsin concentration, extending to a maximum value of ~1650 ng/cm^2^, and did not increase with higher concentrations (*i.e*., 100% SF, non-diluted). Instead, we expected differences between treatments in the dissipative properties of the films. At 1%, 5%, and 50% concentration, the dissipation values for all treatments were similar between them, increasing from ~2×10^−6^ to ~3×10^−6^ and finally to ~7.5×10^−6^. At the highest concentration (100%), however, *d* values were different for each treatment. For nontreated SF, *d* values continued to increase, from 7.5×10^−6^ to 10.0×10^−6^. SF+HAase also presented an increase in dissipation with increasing concentration, from 7.5×10^−6^ to 9×10^−6^. For SF+Trypsin, the increase in *d* was similar to what was observed for SF+HAase, increasing from 7.5×10^−6^ to 9×10^−6^. SI Table 3 summarizies these values. Collectively, *Γ* and *d* suggest that films formed at 1%, 5%, and 50% concentrations are very similar independent of treatment, but not at 100% concentration, despite having similar surface mass densities. These values indicate that the loosely bound molecules adsorbed onto the primary layer are most likely presenting differences in orientation, conformation, and/or grafting densities.

**Figure 2.**
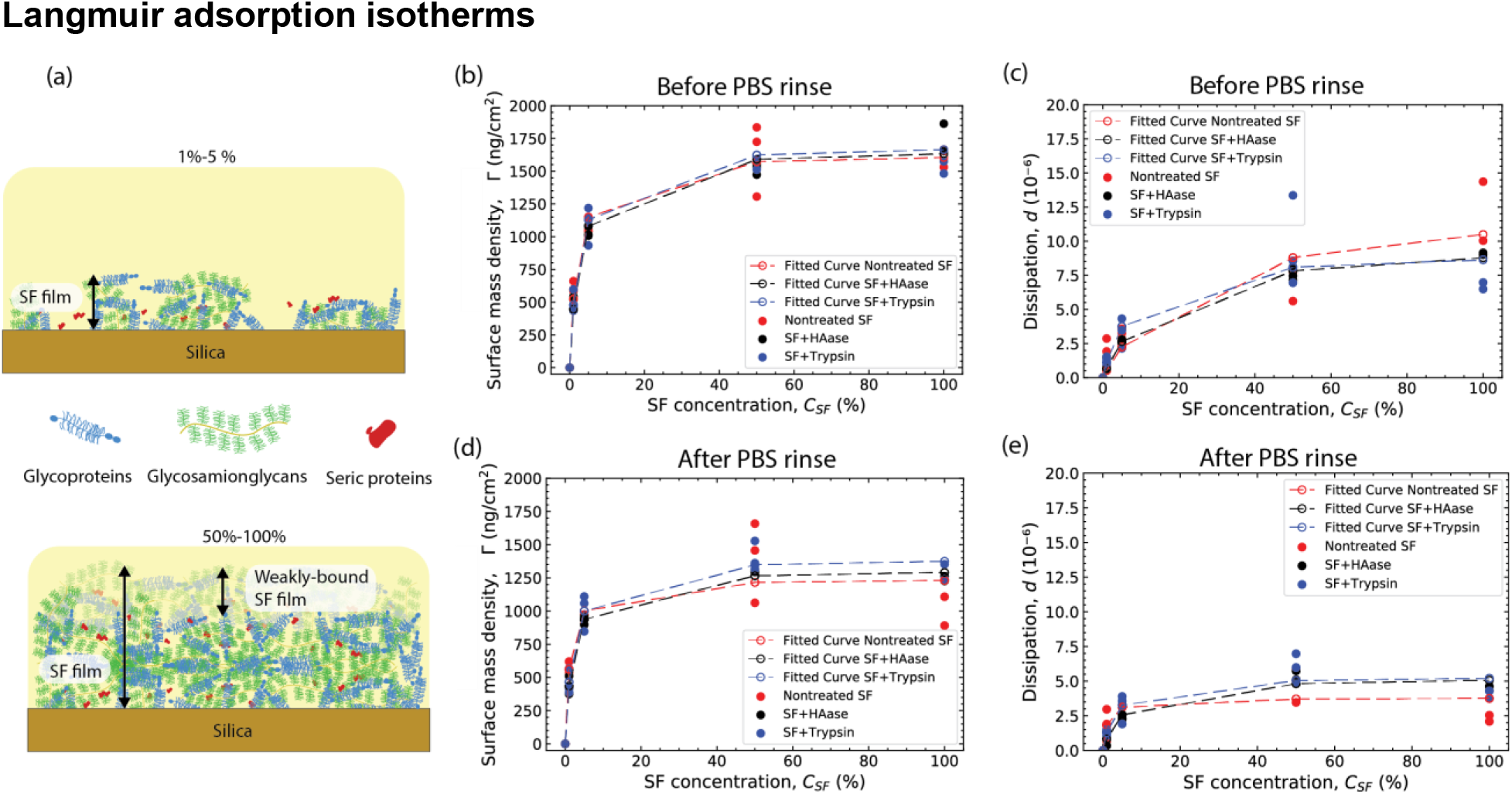
Surface density and dissipation changes with SF concentration. (a) Schematic representation of partially formed and fully formed SF films. (b) Surface density and (c) dissipation of tightly bound films and excess and (d) surface density and (e) dissipation of the tightly bound films left after PBS rinse formed by nontreated SF (red), SF+HAase (black), and SF+Trypsin (blue) before a PBS rinse. Dashed lines represent fitting curves for the Langmuir model, eq. 4.

Next, rinsing with PBS to remove loosely bound molecules revealed a similar qualitative trend to what we observed and described for films before the rinsing step. 1% and 5% dilutions of nontreated SF, SF+HAase, and SF+Trypsin formed partial films with surface mass densities (*Γ*) of ~500 ng/cm^2^ and ~1000 ng/cm^2^, and *d* values of ~2.0×10^−6^ and ~3.0×10^−6^, respectively. A fully formed, tightly bound film was reached with at least 50%nontreated SF, SF+HAase, or SF+Trypsin, with a measured *Γ* of ~1250 ng/cm^2^ and dissipation of ~5.0×10^−6^. At 100% nontreated SF, SF+HAase, and SF+Trypsin, *Γ* did not further increase and remained like what was measured at 50% for all three treatments, ~1250 ng/cm^2^. The dissipation *d*, however, were not similar between treatments. We measured a*d* value of ~5.0×10^−6^ for SF+HAase and SF+Trypsin, and a lower *d* for nontreated SF, ~4.0×10^−6^. The drop in *d* after the PBS rinse is an indication that the film is more rigid, possibly due to a different supramolecular assembly sequence of events, yielding a more compact and dense film, as further discussed in the *Discussion* section.

### Compliance of SF films

For films that are stiffer than the surrounding liquid and considerably thinner than the penetration depth of the shear weave used (~150 nm for the third overtone, *n* = 3), the elastic component of the shear dependent compliance (*ν*) can be obtained from SI eq. 2. Figure 3 shows *d* as a function of *Γ*. In this plot, we remove the symbol delta from the dissipation and surface mass density. These averaged values are not reported as a function of time but rather averaged values at well-defined time points of the measurement (*i.e*., before rinsing step, end of the experiment). SI Figure 3 shows the region from which data points for the elastic component of the compliance was extracted. We identified two regimes for films before rinsing with PBS, Figure 3(a). A less compliant regime, corresponding to the partially formed films (high dilutions, 1% and 5%), and a more compliant regime for the fully formed films (low or no dilution, 50% and 100%). *ν*, obtained from SI eq. 2 and corresponding to the slopes in Figure 3 were 3.2×10^−3^ cm^2^/ng, 2.5×10^−3^ cm^2^/ng, and 2.8×10^−3^ cm^2^/ng for nontreated SF, SF+HAase, and SF+Trypsin for partially formed films, respectively. Fully formed films, at concentrations of 50% and 100% were considerably more compliant, with *ν* values of 7.1×10^−3^ cm^2^/ng, 8.5×10^−3^ cm^2^/ng, and 10.0×10^−3^ cm^2^/ng for nontreated SF, SF+HAase, and SF+Trypsin. From these measurements, we observe that nontreated SF films are less compliant than SF+HAase and SF+Trypsin.

**Figure 3.**
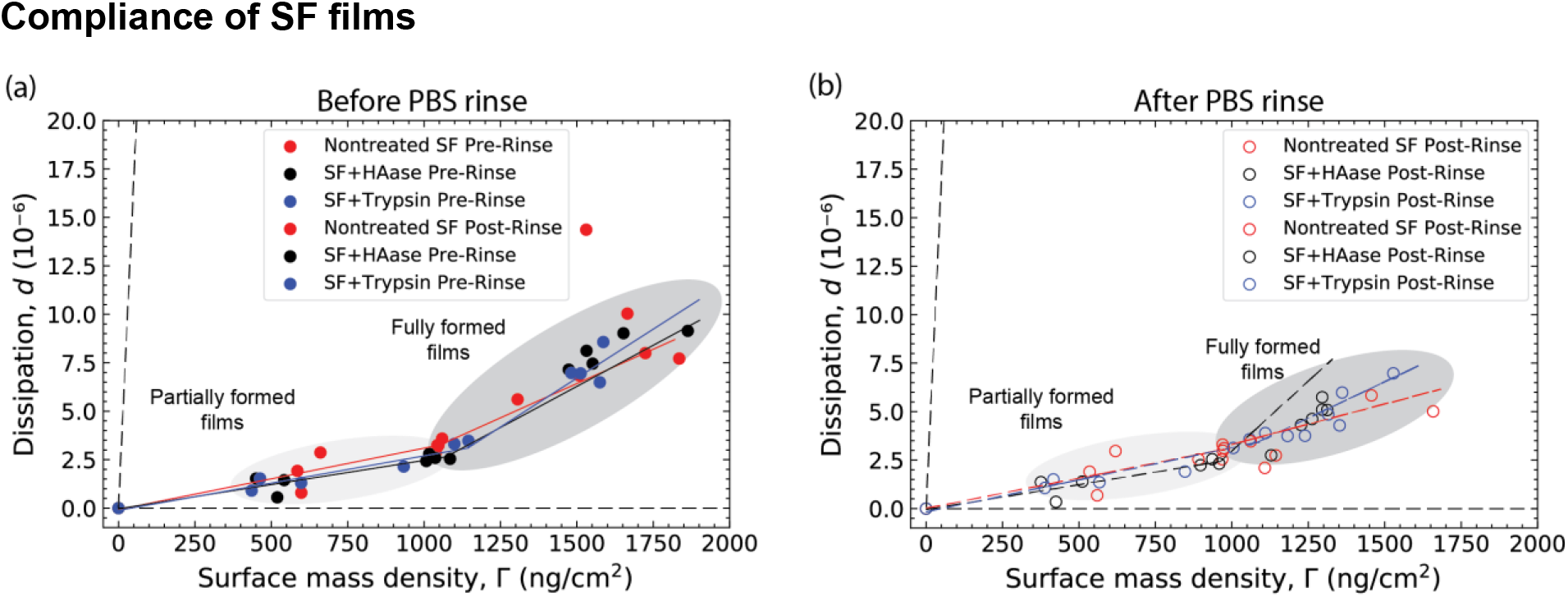
Elastic component of the shear dependent compliance. Change in dissipation as a function of surface density obtained for the adsorption of nontreated SF (red), SF+HAase (black), and SF+Trypsin (blue) for (a) before (filled circles) and (b) after (open circles) a PBS rinse.

Next, we quantified *ν* for films after a PBS rinsing step to remove loosely bound SF components and characterize the compliance of tightly bound films. *ν* remained unchanged for partially formed films (1% and 5%), 3.1×10^−3^ cm^2^/ng, 2.5×10^−3^ cm^2^/ng, and 3.3×10^−3^ cm^2^/ng for nontreated SF, SF+HAase, and SF+Trypsin, respectively. It was for fully formed films that we observed a considerable change in compliance. The rinsing step left behind a less compliant, tightly bound film with *ν* being 4.1×10^−3^ cm^2^/ng, 2.5×10^−3^ cm^2^/ng, and 7.5×10^−3^ cm^2^/ng for nontreated SF, SF+HAase, and SF+Trypsin, respectively. These results suggest that SF fluid films, independently of treatment, form films with a compliance gradient, being less compliant at the substrate interface, and become more compliant at the bulk SF interface. The effect of treatment changes the magnitude of the gradient.

### Normal interaction forces between mica surfaces across undiluted SF

To determine the effect of the enzymatic treatments on the load-bearing properties of SF, we used the SFA to measure normal forces between undiluted (100%) nontreated SF, SF+HAase, or SF+Trypsin. Figure 4(a) shows the experimental schematic and representative force-distance profiles for SFs. Figure 4(b)-(e) and SI Figure 4 displays normal forces, reported as *F/R*, *F* being the normal force and *R* the surface radius of curvature as a function of the surface separation distance, *D*. For all treatments, the force-distance profiles were purely repulsive, but the range and the magnitude of the repulsion depended on the enzymatic treatment and loading/unloading history. Figure 4(b)-(e) compares forces measured for nontreated SF (red curves), SF+HAase (black curves), and SF+Trypsin (blue curves) for subsequent measurements performed on the same location, with 10 min waiting period between measurement. There are qualitative and quantitative differences between force curves. For the first loading/unloading cycle, Figure 4(b), the onset of repulsion, *D*_0_, for all conditions occurs at ~210 nm, equivalent to two SF relaxed films, and increases monotonically with decreasing separation. For untreated SF, at a distance less than ~50 nm, the forces increased sharply, but the substrates never deformed/flattened, reaching a hardwall (HW) at ~40 nm, equivalent to two compressed SF films. During unloading, the force-distance profile followed the trajectory of the loading curve, indicating no hysteresis. For SF+HAase, the repulsion continued to increase monotonically with decreasing separation, until reaching a HW at ~40 nm, like nontreated SF. During unloading, however, forces initially decreased sharply, and transitioned to a monotonic force decay at ~50 nm, showing a significant hysteresis between loading and unloading. For SF+Trypsin, the repulsion increased monotonically with decreasing separation, until reaching a critical thickness, ~90 nm at which the forces increased sharply, followed by a monotonic increase, resembling a sigmoidal curve, reaching a HW at ~60 nm. During unloading, forces decreased sharply, but followed the loading curve trajectory afterwards. The behavior observed for SF+HAase and SF+Trypsin suggests that the films experience a molecular re-arrangement due to confinement, that was not observed for nontreated SF.

**Figure 4.**
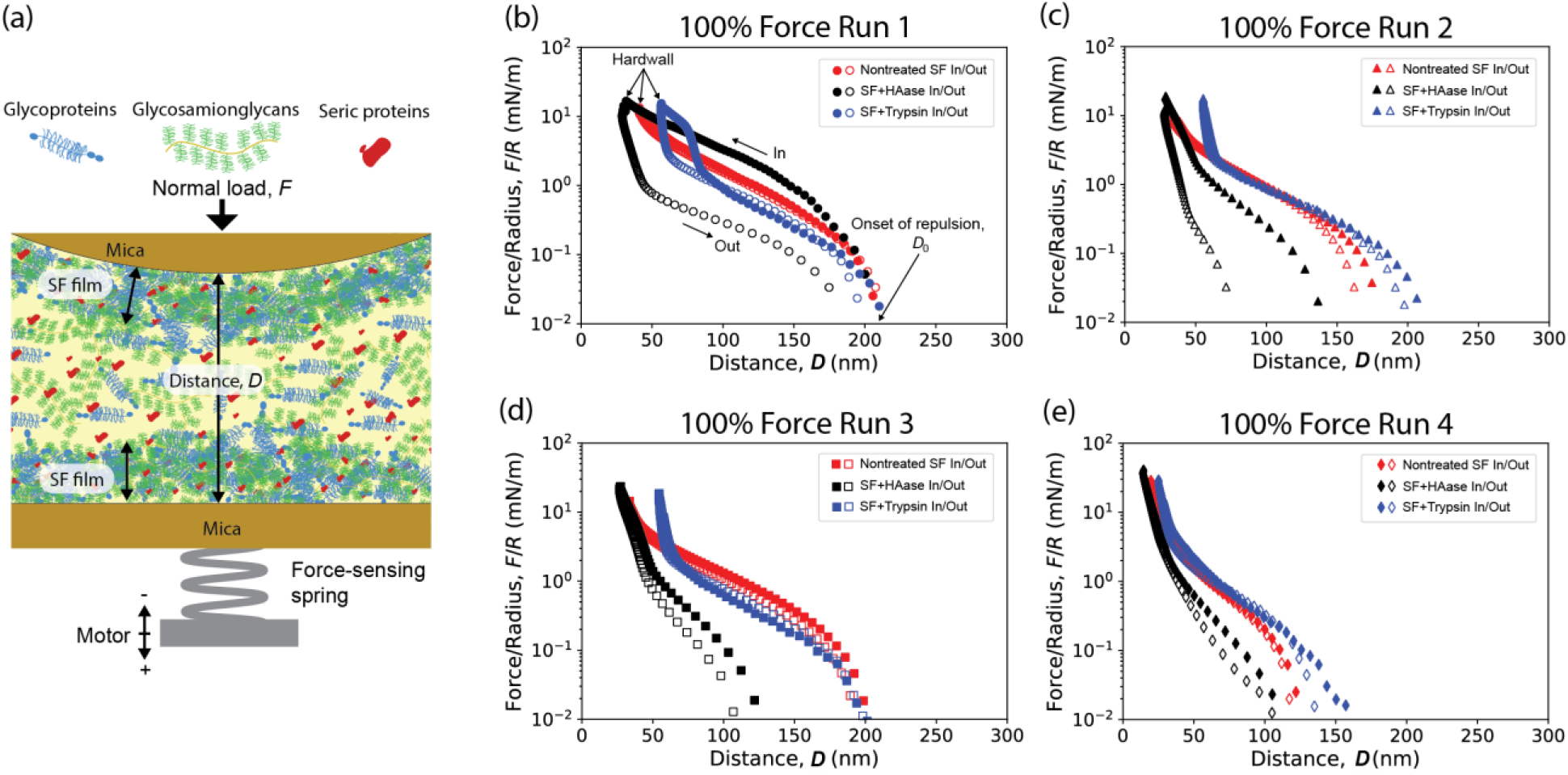
Comparison of normal interaction forces between two mica surfaces across nondiluted SF. All conditions were purely repulsive, and no adhesion was measured. (a) Schematic of the SFA and cartoon representing the main SF components forming a film and confined between mica surfaces. Profiles of the loading (in, filled symbols) and unloading (out, open symbols) for (b) first cycle, (c) second cycle, (d) third cycle, and (e) fourth cycle of nontreated SF, SF+HAase, and SF+Trypsin.

Subsequent loading and unloading cycles in the same location were very similar to the first loading and unloading cycle for nontreated SF, with no change in the onset of repulsion (*D*_0_), hard wall (HW), and unloading, Figure 4(c) and (d) and SI Figure 4(a). That was not the case for SF+HAase, Figure 4 (c) and (d) and SI Figure 4(b). *D*_0_ shifted from ~210 nm to ~150 nm, the force-distance curve of the second compression cycle followed the first unloading force-distance curve very closely, followed by an unloading curve that showed, once again, significant hysteresis. The described behavior repeated for the third loading cycle, with the *D*_0_ shifting form ~150 nm to ~125 nm, however less loading/unloading hysteresis was detected. The substrates never deformed/flattened for this condition. Lastly, for SF+Trypsin, the second and third loading/unloading cycles were almost identical between them, as well as to the unloading force-distance curve of the first compression cycle. That is, *D*_0_ started at ~210 nm increasing monotonically with decreasing separation. At a distance less than ~65 nm, the forces increased sharply, but the substrates never deformed/flattened, reaching a HW at ~60 nm, equivalent to two compressed SF+Trypsin films. The behavior observed for SF+HAase suggests that the films continue to experience molecular re-arrangements due to confinement, which was no longer observed for nontreated SF and SF+Trypsin at similar compression forces and compression rates. Alternatively, it could be possible that by leaving the SF films under an applied load (rather than controlling force) for a finite time, films could experience molecular re-arrangement as well, leading to similar hysteresis. This hypothesis will have to be investigated in future studies.

To test if molecular film re-arrangement continued with higher applied normal forces, we manually compressed (*F/R* > 200 mN/m) nontreated SF, SF+HAase, and SF+Trypsin films until clear flattening occurred, followed by a final motor-controlled loading-unloading cycle, 10 minutes after manual compression. All conditions experienced plastic/viscoelastic deformation, observed by a shift in *D*_0_ to smaller distances. Nontreated SF decreased to ~125 nm, SF+HAase decreased to ~100 nm, and SF+Trypsin decreased to ~150 nm, Figure 4(e). HW values decreased to ~50% of the initially measured HW (FR 1), to ~20 nm, ~25 nm, and ~40 nm for nontreated SF, SF+HAase, and SF+Trypsin, respectively.

To understand the origin of the repulsive interactions described above, “In” force-distance profiles (filled symbols) shown in Figure 4 were fitted with the Alexander-de Gennes (AdG) model, SI eq. 5. Experimental data and fits are shown in SI Figure 5 and fitted parameters summarized in SI Table 4. Our results confirmed that normal forces between two mica surfaces across nontreated SF can be well described by the AdG model, as previously reported.[6,39] For this model, the average effective brush thickness *L* = *D*_0_/2, corresponding to a brush on each mica surface. The average effective brush thickness (*L*) and average effective grafting distance (*s*) values were 93 ± 6 nm and 0.6 ± 0.01 nm, respectively. Based on the coefficient of determination (*r*^2^) from our non-linear least-squares regression of the AdG model, we observed that the prediction as a brush decreases for SF+HAase (*r*^2^ = 0.95 ± 0.02) and SF+Trypsin (*r*^2^ = 0.88 ± 0.07) when compared against nontreated SF (*r*^2^ = 0.98 ± 0.01). The average *L* and *s* values increase to 125 ± 23 nm and 0.47 ± 0.08 nm for SF+HAase, and 107 ± 14 nm 0.54 ± 0.07 nm, for SF+Trypsin, respectively. These results, combined with the force-distance profile shapes, clearly demonstrate that nontreated SF, SF+HAase, and SF+Trypsin form films on silica that respond differently to confinement (compression) due to differences in the supramolecular assembly.

To further describe the repulsive behavior of mica across nontreated SF, SF+HAase, and SF+Trypsin, we additionally used the Dolan and Edwards (DnE) model, SI eq. 6. Experimental data and fits are shown in SI Figure 5 and summarized in SI Table 4 as well. The fitted decay length, *R*_eff_, increased as follows: SF+Trypsin (9.5 ± 8.0 nm), followed by nontreated SF (19 ± 5.0 nm), and SF+HAase (663 ± 1100 nm). The pre-factor *A*, which quantifies the amplitude of the repulsion, varied considerably between treatments, an indicator of the load-bearing capability of the films, and increased as follows: Nontreated SF (84 ± 26 nm), followed by SF+HAase (1320 ± 1070 nm), and lastly SF+Trypsin (1,228,045 ± 1,087,503 nm).

### Normal interaction forces between mica surfaces across diluted (5%) SF

Next, based on the Langmuir adsorption isotherm results, we asked how partially formed films would respond to confinement. We investigated the interaction forces of mica across 5% nontreated SF, SF+HAase, and SF+Trypsin using the SFA. While little to no hysteresis was observed for 5% nontreated SF and for 5% SF+HAase either between in (loading) and out (unloading) or between cycles (Force Run 1-3), 5% SF+Trypsin showed considerable hysteresis, between in/out (loading/unloading) and between cycles (Force Runs). Interestingly, *D*_0_ for Force Run 2 and 3 were very close to the termination of interaction of Force Run 1 and 2, respectively. This observation suggests that the film has undergone long-term molecular rearrangement, SI Fiugre 5 and SI Figure 6.

Similar to what performed for 100% and based on the coefficient of determination (*r*^2^) from our least-squares regression of the AdG and DnE models, summarized in SI Table 5, we observe that the prediction of the confinement force normalized by the radius of curvature as a function of the separation distance is more accurately described as an end-grafted polymer in the mushroom regime for films formed with diluted (5%) SF solutions.

## DISCUSSION

The main goal of this study was to examine the effects of two enzymatic treatments on the formation and shear-dependent compliance of SF nano-films, and interaction forces of macroscopic surfaces across SF adsorbed to oxide surfaces (e.g., silica and mica). SF components known to contribute to the load-bearing, wear protection, and lubrication of articular cartilage surfaces were digested using GAG and protein specific enzymes. We used HAase to depolymerize HA. However, it is very likely that other GAGs, such as chondroitin sulfate were also degraded, given the structural similarity. In fact, it has been reported that HAases have strong hydrolytic activity toward chondroitin sulfate comparable to that for HA.[40] To cleave SF proteins, we used trypsin, a protease that reacts with peptide bonds between carboxylic acid groups of lysine or arginine and the amino group of the adjacent amino acid residue, with its efficacy depending on the tertiary and quaternary protein structure, as some of the peptide bonds may become inaccessible.[41] This treatment is effective on large glycoproteins, such as lubricin[7,30] but will also target smaller glyco- and globular proteins, like decorin and albumin, respectively, however less efficient on the latter given its compact structure. Osteo- and rheumatoid-arthritic SFs have been show to present, with varying concentrations, enzymes such as cathepsin,[42,43] plasmin,[44,45] and higher HAase activity compared to that of healthy SFs.[46] Cartilage from human femoral head has a reported surface charge density of 37 mC/m^2^,[47] which is within range to that for silicates, 3.2-80 mC/m^2^ used in this study.[48,49] Therefore, findings reported here can provide insights into the formation of films by healthy (nontreated SF) or pathophysiologic SFs (SF+HAase or SF+Trypsin) on either implanted surfaces (e.g., oxides) or the reformation of torn lamina splendens on the articular cartilage surface.

At low concentrations (1-5%), film-formation kinetics results revealed that only trypsin altered the characteristic saturation time, decreasing it by a factor of two, SI Figure 2 and SI Table 1. This could be due to the smaller radius of gyration of the peptide fragments combined with the positively charged ends (arginine and lysine), leading to a faster migration and adsorption to the negatively charged silica and mica surfaces. Adhesion studies of small molecules having cationic amino acids, in particular lysine, have been proposed to drive electrostatic adsorption to mica surfaces in high ionic strength environments (*I* ≈ 150 mM).[50,51] The fact that the saturation times for nontreated SF and SF+HAase are similar suggests that glycoproteins, such as lubricin, which carry two positively charged globular end-domains, could be driving the adsorption process. At higher concentrations, however, the main adsorption mechanisms are dominated by bulk transport, as the saturation time for all treatments tested are similar. Bulk transport has been observed for human serum albumin studies adsorbed to silica resonators using QCM-D at concentrations 5-10 mg/ml, equivalent to the concentrations found in SF (our 50% and 100% dilutions, respectively).[39,52] However, given the complex composition of SF, different processes such as desorption/adsorption, competitive exchange, and exchange via transient complex formation will also participate in the film formation and maintenance of the film.

SF is known to be rheopectic at low shear rates but shear-thinning at high shear rates.[6,53] We observed rheopectic behavior for nontreated SF and SF+HAase, but not for SF+Trypsin, for which we qunatified shear-thinning response only, SI Figure 8. We also quantified the bulk refractive index of SFs and did not find a correlation between refractive index and treatment, SI Figure 9.

From the Langmuir adsorption isotherms, Figure 2, we observed that full film formation is achieved with at least 50% SF concentrations for all treatments. This observation could be interpreted as a protection mechanism to ensure instantaneous surface saturation to reform a protecting film, even if local concentration variations are present due to, for example, large depletion forces driven by external macroscopic factors (*e.g*., locomotion).[54] Furthermore, this film could be an essential precursor step for the formation of the lamina splendens,[6,7] which has been implicated in providing wear protection and lubrication,[6,55,56] as well as regulating the permeability and compression resistance of the superficial layers of cartilage.[57] Films, regardless of treatment and concentration, consist of a strongly adsorbed layer, followed by a second loosely bound layer that could be easily removed by rinsing with PBS. The pre-rinse and post-rinse films showed mechanical differences between them, Figure 3, most likely because of structural, compositional, and hydration differences. Pre-rinse films did not reveal apparent differences between treatments in their shear-dependent compliance, but confinement showed significant differences in the interaction forces, Figure 4. A mechanical analogy describing the film responses during confinement is as follows: for nontreated SF films, we observed a dominant elastic response (for the investigated loading/unloading rates and forces applied), which could be modeled as a series of parallel one-dimensional springs. In our configuration, the film dimensions (*L* ≈ 100 nm) are considerably thinner than the contact diameter (ϴ > 10 μm). That is, the film thicknesses are at least two orders of magnitude smaller than the contact diameter. For these conditions, the multi-spring model, the Winkler model, can be used. Other studies have used it to approximate the compression forces of hydrogels[58] and protein films.[59] The springs have a specified compression range, corresponding to the indentation. For a compression spring, the stiffness is determined by the number of active coils per unit length, the diameter of the coil, and the diameter of the wire. SF+Trypsin films can be modeled with the same spring (*i.e*., identical coils per unit length, coil diameters, and wire diameters) as the nontreated SF films, however, with a shorter compression range, as the compression force-distance initial slope is very similar as the nontreated SF until approximately 65 nm, were the force rapidly increases. This response would correspond to the bottoming or full compression of the spring. We propose to describe the compression of SF+HAase films as a series of dashpots in parallel, as the force-distance profiles showed a significant hysteresis with no recovery during experimental time scales, which could be due to the molecular re-arrangements leading to enhanced hydrophobic interactions favored by confinement, overcoming entropic stabilization driven by excluded volume. The mechanical analogies for the films are schematized in Figure 6. The AdG and DnE model described well the interaction forces of mica surfaces across SFs, and suggest that the elastic films (nontreated SF) are better modeled as fully swollen polymer brushes and that enzyme treated SFs films (SF+HAase and SF+Trypsin) are better modeled as polymer mushrooms.

**Figure 6.**
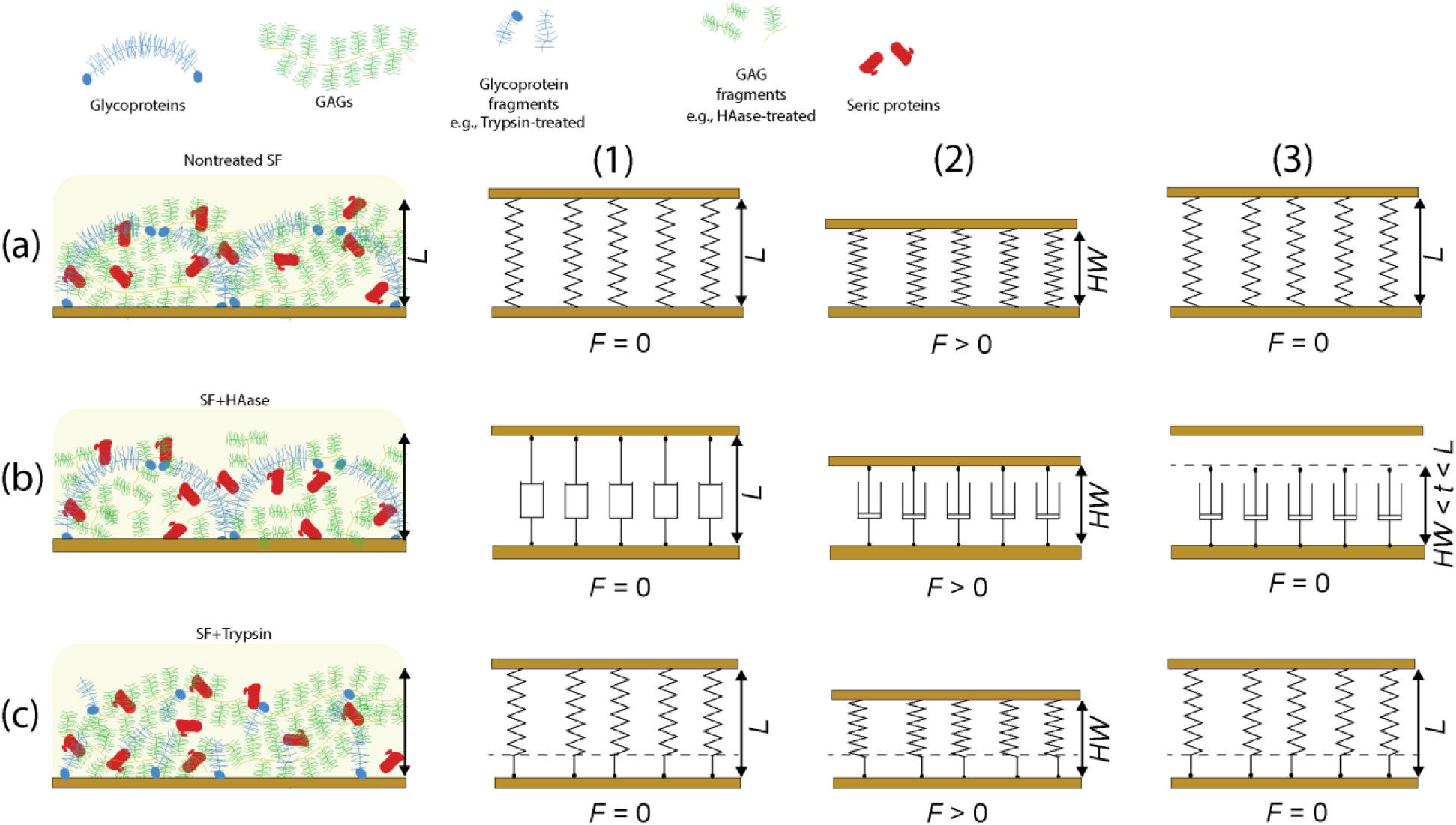
Mechanical model for SF films. (a) nontreated SF with the mechanical spring model – long spring, (b) SF+HAase with the mechanical dashpot model, and (c) SF+Trypsin with the mechanical spring model – short spring. (1) shows the initial “in rest” configuration of the SF films, where the thickness is equivalent to *D*_0_. (2) represents the maximum applied force configuration, reaching a film thickness equivalent to the *HW*. (3) corresponds to the configuration of the films after removing the applied load.

It is interesting to note that HA and its complexes provided stiffness to the tested films under confinement, while lubricin and other glycoproteins provided an extended elastic range. It is well documented that HA regulates the viscosity of SF in bulk, and mediates elastoviscous lubrication.[21] On the other hand, lubricin is now accepted as the main molecule mediating boundary lubrication.[14] However, building body of evidence suggests that the synergistic surface interactions of SF components are responsible for the extremely efficient wear protection and lubrication properties of articular cartilage surfaces,[19,23,27,29] and that the film can strongly affect the rheological an tribological behaviors of viscosuplements.[19,21,60] Based on these reports, it is crucial to understand and identify the precursor films’ properties (e.g., chemical make-up, stiffness, roughness, etc.).

Elasticity and robustness of nontreated SF films is desired. For example, synovial joint situations in which the tangential velocity is very small or zero (the investigated condition) are “heel-strike” or “toe-off”. In the absence of relative motion between the interacting surfaces (*e.g*., apposing cartilage surfaces), hydrodynamic or elasto-hydrodynamic lubrication cannot exist, and the surfaces experience pure normal loads. A study in the 50’s reported that SF has a special capacity for forming a layer with some degree of elasticity.[5] Here, we report that both biomolecules (or more broadly, GAGs and proteins in SF) are crucial for the proper load-bearing properties of films, and that nontreated SF films have some degree of elasticity indeed.

Furthermore, we report that by digesting GAGs or proteins, the elasticity is removed or reduced, respectively. These findings emphasize the importance of the synergistic interactions of SF components not only for the lubrication and wear protection of the surfaces (natural or synthetic), as reported extensively, but also for the proper load-bearing properties of the film. Indeed, it can be the poor load-bearing capabilities of the films that lead to improper lubrication. Even more, the combined effects could then lead to excessive interstitial fluid pressurization,[61,62] essential to the load-bearing and lubrication properties of articular cartilage. While effects of loading/unloading rates, contact times, contact loads, are all important parameters to fully understand the mechanisms of SF films under confinement and the effect of enzymatic treatments, it is beyond the focus of this study and is being systematically investigated.

Oxides such as alumina and ultra-high molecular weight polyethylene (UHMWPE) are frequently used for artificial joint surfaces, due to their mechanical properties, biocompatibility, and good tribological performance. A well-known failure mode in artificial joints is due to the release of UHMWPE wear particles leading to abrasion or release of metal and metal oxide ions due to tribocorrosion, both activating adverse biological responses.[63–65] Surface functionalization approaches, such as grafting of polymer brushes or plasma treatments to modify surface charge and control adsorption of SF components has been intensively investigated in the last decade, with the goal of enhancing the tribological performance of the interacting implant surfaces.[66,67] By controlling the adsorption of SF components, it could become possible to tune the tribolayer that forms at the surface of implanted joints. These layers have been suggested to improve load distribution imposed by the orthopedic implant.[68] If the initially formed films on implant surfaces is built by pathological SF, as modeled here by enzymatic treatments, it is clear that the load distribution of the films would be not as efficient as that of films formed by healthy SF, potentially leading to a poor tribolayer formation. Given that implants, such as total hip or knee replacements are commonly placed in patients that have synovial joint pathophysiology, understanding how SF components interact with surfaces is of crucial interest to identify the suitable surface functionalization strategies that yield SF films with the desired physico-chemical properties.

## CONCLUSIONS

Collectively, our results suggest that SF forms fully saturated films at concentrations of at least 50% of what is found at physiological concentrations, and that the removal of the excess layer exposes differences in the shear dependent elastic component of compliance of the different SF films. Furthermore, we find that the absence of either GAGs or proteins abolishes the load-bearing capabilities, and that the loading curves can be well described by two simple models from polymer physics used to describe the interaction forces of endgrafted polymer chains, de Alexander-de Gennes (AdG) and the Dolan and Edwards (DnE) models. The findings reported can be of relevance to understanding the formation of protecting films on cartilage or implant surfaces and emphasize the importance of load-bearing properties of these SFs under confinement in addition to the tribological characterization, to fully understand the supramolecular assemblies and synergistic interactions of SF components.

## SUPPLEMENTAL INFORMATION

The Python codes written and data files used to generate the figures in this report can be found at: https://gitlab.com/randresen/glycosaminoglycans-and-glycoproteins-influence-the-elastic-response-of-synovial-fluid-nanofilms-on-model-oxide-surfaces

## ACKNOWLEDGMENTS

R.C.A.E, A.S.M., and A.M.S. acknowledge funding from the NSF-CREST:Center for Cellular and Biomolecular Machines through the support of the National Science Foundation (NSF) Grant No. NSF-HRD-1547848). R.C.A.E. acknowledges funding from the NASA Merced nAnomaterials Center for Energy and Sensing (MACES) through the support of the National Aeronautics and Space Administration (NASA) Grant No. NNX15AQ01. A.M.S. and A.G. acknowledge funding from the career grant NSF CBET 2047210 awarded to A.G.

