## Supporting Information for "Glycosaminoglycans and glycoproteins influence the elastic response of synovial fluid nanofilms on model oxide surfaces"

### EXPERIMENTAL SECTION

#### Quartz Crystal Microbalance with Dissipation (QCM-D)

QCM-D measurements were performed using an openQCM-Q1 microbalance (Novaetech SRL, Pompey, Italy), AT-cut crystals coated with 50 nm silica, 5 MHz fundamental frequency (NanoScience Instruments, QSX 303), and the 3rd overtone ($f_3$). First, silica crystals were cleaned with 2% w/v sodium dodecyl sulfate (SDS) solutions (MP Biomedicals LLC, 811032), followed by 2% w/v digestive enzyme solution (Contrex EZ, 5405), copious amounts of ultrapure water, 70% ethanol, then dried under a stream of compressed nitrogen, and ending with a plasma cleaning step for 2 minutes (PDC32-G, Harrick, USA) before use. The baseline for the measurements were obtained by flowing PBS through the system and allowing for equilibration, using a peristaltic pump set at ~75 µL/min (Golander LLC, BQ80S Microflow Variable-Speed). SF solutions (nontreated SF, SF+HAase, or SF+Trypsin) were then pumped through the system, filling the fluid cell and stopping the flow. After surfaces reached saturation, approximately after 30 minutes, PBS was pumped to remove loosely bound molecules, referred to as rinsing step. After each experiment, the tubing and system were cleaned with 2% w/v digestive enzyme solution, 2% w/v SDS, and ultrapure water. All measurements were conducted at 25 °C.

The resonance frequency ($f$) of a quartz crystal microbalance in contact with liquid is determined as described by Kanazawa and Gordon.[1,2] The dissipation ($d$) of the crystal for our system is determined by measuring the full with half max (FWHM) of the resonant conductance curve.[3,4]

***Theoretical modeling of the QCM-D data.*** In QCM-D measurements, a quartz crystal oscillator is set to oscillate at its resonance frequency. The shift in frequency due to the formation of an SF adsorption layer is typically 10-200 Hz. The frequency and dissipation changes can be related to the mass oscillating with the crystal and the viscoelastic properties of the layer through various models.[5]

***The Sauerbrey model.*** The Sauerbrey model, equation 1, relates the frequency change ($\Delta f_n$) of the oscillating quartz crystal, due to the presence of a thin film, to the change in surface mass density of the film ($\Delta \Gamma$):[6]

$$\frac{\Delta f_n}{n} = -\frac{f_0}{t_q \rho_q}\Delta\Gamma = -c\Delta\Gamma \quad (1)$$

where $n$, $f_0$, $t_q$, $\rho_q$ are the overtone number, the fundamental frequency of the quartz oscillator, the thickness, and the density of quartz, respectively. The value for $c$ of the crystals used is 17.7 ng $Hz^{-1}$ $cm^{-2}$.[Citation error] The Sauerbrey equation gives the surface mass density, including liquid inside the film. Eq. 1 is valid only for thin films in air. However, for lateral homogeneous films with a change in dissipation value larger than zero ($\Delta d$ > 0), the elastic component of the shear dependent compliance ($\nu$) can be approximated from the ratio between the change in dissipation ($\Delta d$) and the change in surface mass density ($\Delta\Gamma$):[5,7,8]

$$\nu = -\frac{\Delta d}{\Delta\Gamma} \quad (2)$$

In QCM-D, the dissipation is defined as the sum of all energy losses in the system per oscillation cycle, and it is a dimensionless number, as it relates the energy lost per cycle over the energy stored. An ideally rigid film will have zero dissipation for any change in surface mass density. For a purely viscous film, the maximum possible change in dissipation is -2 for every Hz, which adjusted to surface mass density is 644 $\Delta d$ $ng^{-1}$ $cm^2$. A kinetic model was used to determine the time it took each SF treatment and dilution to saturate the silica surfaces, the saturation time $\tau$, obtained by:[9]

$$\Gamma = \Delta\Gamma_{max}\left(1 - e^{-\frac{t}{\tau}}\right) \quad (3)$$

where $\Gamma$ is the instantaneous surface mass density, $\Delta\Gamma_{max}$ is the maximum measured change in surface mass density, and $t$ the experimental time. The saturation time, $\tau$ was used as free fitting parameter. Time t = 0 corresponds to the injection of either nontreated SF, SF+HAase, or SF+Trypsin.

***The Langmuir isotherm model.*** The Langmuir model, equation 4, describes the coverage of molecules adsorbed on a solid surface:[9]

$$\Gamma = \frac{\Gamma_{max} K_a [SF]}{1 + K_a [SF]} \quad (4)$$

where $\Delta\Gamma$ is the change in surface mass density, $\Delta\Gamma_{max}$ is the change in surface mass upon saturation, and $K_a$ is the equilibrium constant. While this model is valid for single components, it will be used to describe the coverage of SF molecules adsorbed on the silica surface at different dilutions, with the understanding that SF is a multicomponent and complex fluid.

**Surface Forces Apparatus (SFA)**

***SFA surface preparation.*** SFA experiments were performed using an SFA 2000 (SurForce LLC, CA, USA). Surfaces were prepared using established procedures.[10,11] Two back silvered mica surfaces, 2–3 µm thick (S&J Trading INC, ruby mica grade #1 V-1/V-2), were glued on cylindrical glass discs with a radius of curvature of $R$ = 2 cm using thermosetting glue (EPON, 1004F). After gluing the mica surfaces, one surface was mounted on a double cantilever spring, with an optically calibrated spring constant of $k_N$ = 471 N/m, resulting in a force (normalized by the radius of curvature, F/R) resolution of ~0.001 mN/m. The other mica surface was installed on a top holder, facing the lower surface in a crossed-cylinder configuration, building an optical cavity. Using Multiple Beam Interferometry, the thickness of the mica surfaces was measured at contact in air.[12] The wavelength of the fringes of equal chromatic order (FECO) were recorded using a scientific digital camera (Hamamatsu Orca $R^2$, Japan) and post analyzed using an in-house MATLAB (MathWorks, R2020b) script. After the calibration step, surfaces were adjusted to a separation distance (*D*) of ~1 mm and 30 µL of nontreated SF, SF+HAase, or SF+Trypsin injected between the surfaces. The bottom of the SFA chamber was filled with water to saturate the atmosphere with water vapor and minimize the evaporation of SF during the experiment.

***SFA measurements.*** SFA measurements of the normal interaction forces (*F*) were performed using well-established procedures.[10] After letting the SF solutions saturate the surfaces for 1 hr., surfaces were brought together manually to a separation distance of ~1 µm. Then, surfaces were approached (in) or separated (out) at a constant velocity of ~15 nm/s using a DC motor with a 1670:1 gear ratio (Faulhaber Micromo). The separation distance (*D*) between the surfaces and the normal interaction force were measured using FECO analysis.[12] All experiments were performed at 25 °C. Three consecutive measurements were performed in each new position (FR1-3), with 10 min intervals between compression (in/out) cycles. This was followed by a manual-driven compression resulting in surface flattening, finishing with a last motor-controlled *F-D* measurement (FR4). Two positions per sample were tested.

***The Alexander-de Gennes model (AdG).*** The AdG model, equation 5, relates the force normalized by the radius of curvature (*F/R*) of the interacting films to the equilibrium film thickness (*L*) and the average spacing between close neighboring chains (*s*) of a polymer brush:[13]

$$\frac{F}{R} = \frac{16\pi kTL}{35s^3}\left[7\left(\frac{2L}{D}\right)^{5/4} + 5\left(\frac{D}{2L}\right)^{7/4} - 12\right] \quad \text{for } D < 2L \tag{5}$$

where *k* is the Boltzmann constant, and *T* is the experimental temperature. The first term comes from the osmotic pressure, which increases as the two surfaces approach each other. The second term accounts for the decrease in elastic energy as the films are compressed. This model was originally developed to describe the interaction forces of neutral, grafted polymer brushes. This work does not necessarily imply that the SF films would adopt a well-defined dense brush conformation on the mica substrate. It instead suggests that the film can be described as a purely repulsive effective brush layer. The use of this model is justified by the high salinity of the medium (PBS ~150 mM, with Debye length, $\kappa < 1$ nm), which is expected to screen most of the electrostatic interactions and can therefore be considered neutral.

***The Dolan and Edwards model (DnE).*** The DnE model, equation 6, relates the force normalized by the radius of curvature (*F/R*) of the interacting films to the effective coil size ($R_{eff}$) and the area per chain (*a*):[14]

$$\frac{F}{R} = \frac{72\pi kT}{a} e^{-\frac{D}{R_{eff}}} \quad \text{for } D < R_{eff} \tag{6}$$

where *k* is the Boltzmann constant, and *T* is the experimental temperature. This model was developed to describe the interaction forces between two surfaces with sparse, end-grafted chains in a theta solvent, that is, engrafted polymers adopting a mushroom configuration, instead of a well swollen polymer brush (AdG). This work does not necessarily imply that the SF films would adopt a mushroom conformation on the mica substrate, instead, it suggests that the film can be described as a purely repulsive mushroom layer. In this manuscript we report the pre-factor $A = \frac{72\pi kT}{a}$, which provides a scaling for the amplitude of the repulsion.

**Bulk rheometry.** To test the effects of enzymatic treatment on SF, we performed bulk rheology measurements. An Anthon-Paar rheometer (model MCR-102) with parallel stainless-steel plate attachments (Anthon-Paar, PP25, diameter 25 mm) was used to probe the SF sample. Data was collected using Rheocompass. A working gap of 500 µm was used in all experiments. Tests were performed at room temperature (25 °C) to confirm that enzymatic treatments, SF+HAase and SF+Trypsin, altered the bulk viscoelastic properties of SF. Nontreated SF was used as a control. Viscosity as a function of shear rate is shown in SI Figure 8, for nontreated SF, SF+HAase, and SF+Trypsin. 300µL total volume of each SF treatment were loaded to the bottom parallel plate of the rheometer. The top plate was lowered until the final gap reached 500µm; the assembly was then left for 5 minutes to equilibrate. Viscosity measurements were taken at constant flow for 5 minutes at a specific shear rate. Shear rate was varied from 0.01 to 1000 1/s.

**Refractive index measurements.** A benchtop Abbe refractometer (Atago Co.) was used to determine the refractive index ($\eta$) of nontreated SF, SF+HAase, and SF+Trypsin with ± 0.0005 resolution.

### RESULTS

#### Film formation kinetics

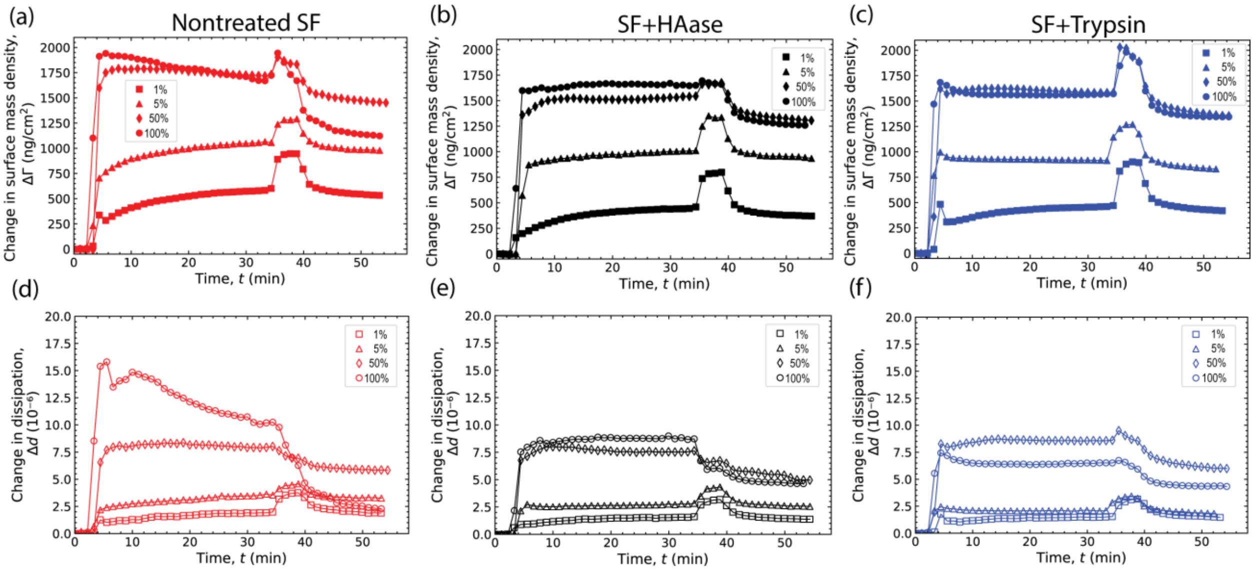


**SI Figure 1. Change in surface mass density and change in dissipation during the adsorption of SFs to silica.** (a) Nontreated SF, (b) SF+HAase, and (c) SF+Trypsin at the four tested concnetrations, 1%, 5%, 50%, and 100%. For clarity, only every 100th data point is shown.

**SI Table 1. Film formation kinetics parameters.** Saturation time, $\tau$, from eq. 3, and surface mass density lost during the PBS rinse, $\Delta\Delta\Gamma$.

| Condition | $\tau_{1\%}$ (min) | $\Delta\Delta\Gamma_{1\%}$ (ng/cm$^2$) | $\tau_{5\%}$ (min) | $\Delta\Delta\Gamma_{5\%}$ (ng/cm$^2$) | $\tau_{50\%}$ (min) | $\Delta\Delta\Gamma_{50\%}$ (ng/cm$^2$) | $\tau_{100\%}$ (min) | $\Delta\Delta\Gamma_{100\%}$ (ng/cm$^2$) |
|---|---|---|---|---|---|---|---|---|

| | | | | | | | | |
|---|---|---|---|---|---|---|---|---|
| **Nontreated SF** | 3.4 ± 1.9 | -44 ± 6 | 2.0 ± 1.1 | -76 ± 11 | 0.9 ± 0.3 | -230 ± 47 | 0.7 ± 0.3 | -520 ± 142 |
| **SF + HAase** | 4.0 ± 1.9 | -67 ± 32 | 2.1 ± 0.9 | -105 ± 23 | 0.9 ± 0.2 | -218 ± 35 | 0.9 ± 0.3 | -532 ± 128 |
| **SF + Trypsin** | 1.6 ± 0.9 | -42 ± 9.5 | 0.7 ± 0.1 | -95 ± 13 | 0.9 ± 0.5 | -331 ± 207 | 1.0 ± 0.2 | -320 ± 151 |

### Langmuir adsorption isotherms

**SI Table 2. Langmuir model fitting results.** Maximum surface mass density ($\Gamma$), maximum dissipation ($d$), and equilibrium constant ($K_a$) values obtained by using the Langmuir model, eq. 4.

| **Condition** | **Maximum surface mass density, $\Gamma_{max}$ (ng/cm$^2$)** | | **Maximum dissipation, $d_{max}$ (x10$^{-6}$)** | | **$K_a$ (min)** | |
|---|---|---|---|---|---|---|
| | **Pre-rinse** | **Post-rinse** | **Pre-rinse** | **Post-rinse** | **Pre-rinse** | **Post-rinse** |
| Nontreated SF | 1638.8 | 1245.5 | 13.0 | 3.8 | 0.47 | 0.82 |
| SF+HAase | 1678.8 | 1317.0 | 10.0 | 5.3 | 0.36 | 0.49 |
| SF+Trypsin | 1707 | 1402.5 | 9.3 | 5.4 | 0.39 | 0.50 |

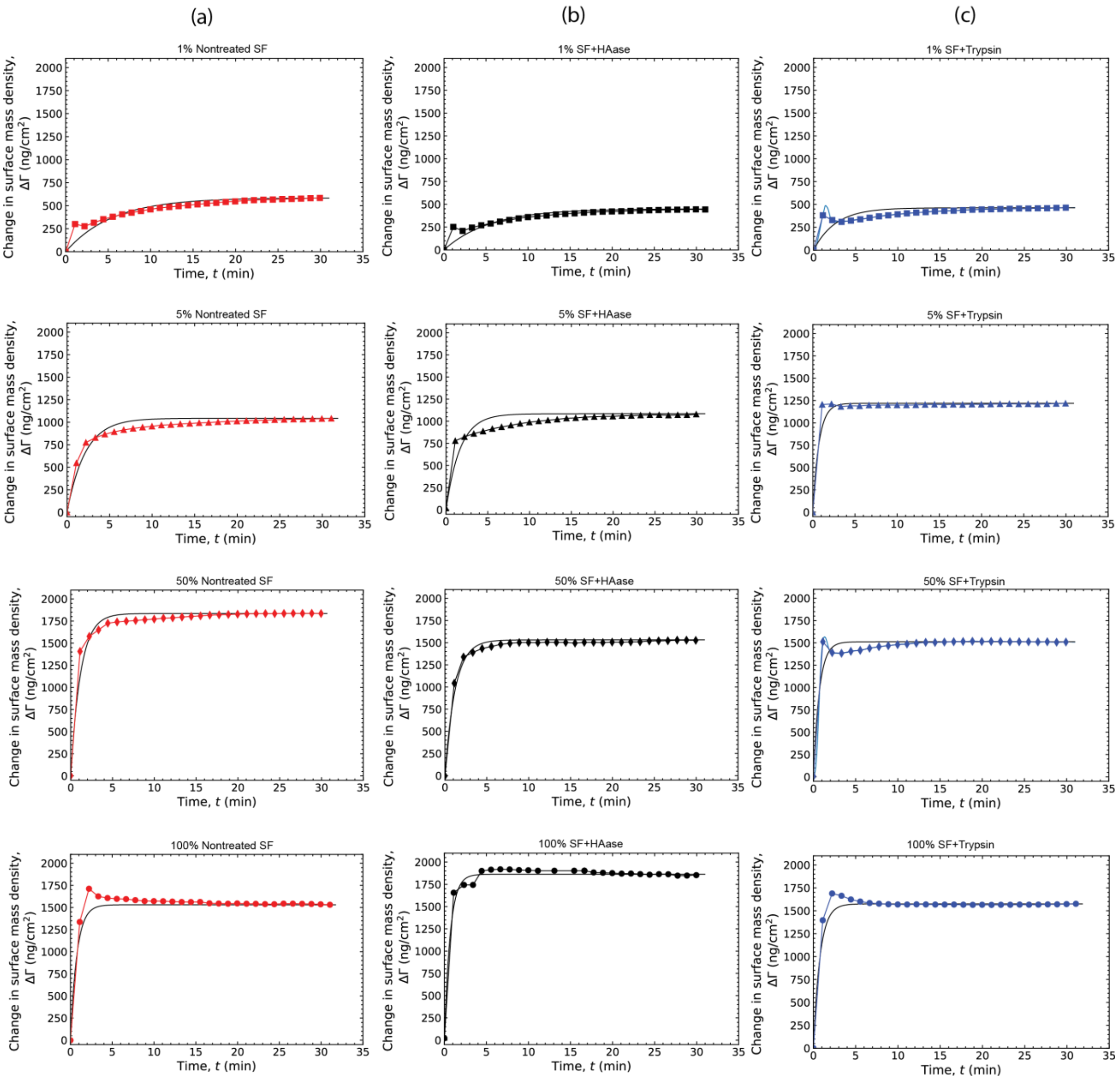


**SI Figure 2. Change in surface mass density during the adsorption of SF to silica.** (a) Nontreated SF, (b) SF+HAase, and (c) SF+Trypsin. For clarity, only every 100th data point is shown.

**SI Table 3. Langmuir model fitting results.** Maximum surface mass density ($\Gamma$), maximum dissipation ($d$), and equilibrium constant ($K_a$) values obtained by using the Langmuir model, eq. 4.

| **Condition** | **Maximum surface mass density, $\Gamma_{max}$ (ng/cm$^2$)** | | **Maximum dissipation, $d_{max}$ (x10$^{-6}$)** | | **$K_a$ (min)** | |
|---|---|---|---|---|---|---|
| | **Pre-rinse** | **Post-rinse** | **Pre-rinse** | **Post-rinse** | **Pre-rinse** | **Post-rinse** |
| Nontreated SF | 1638.8 | 1245.5 | 13.0 | 3.8 | 0.47 | 0.82 |
| SF+HAase | 1678.8 | 1317.0 | 10.0 | 5.3 | 0.36 | 0.49 |
| SF+Trypsin | 1707 | 1402.5 | 9.3 | 5.4 | 0.39 | 0.50 |

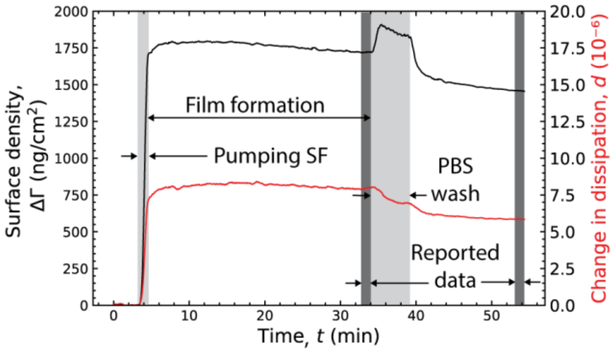


**SI Figure 3. QCM-D experimental timeline.** Surface density as a function of time indicating the pumping of 50% nontreated SF, the film formation, PBS wash, and the regions used to collect reported data for intrinsic viscosity.

### Normal interaction forces between mica surfaces across undiluted SF

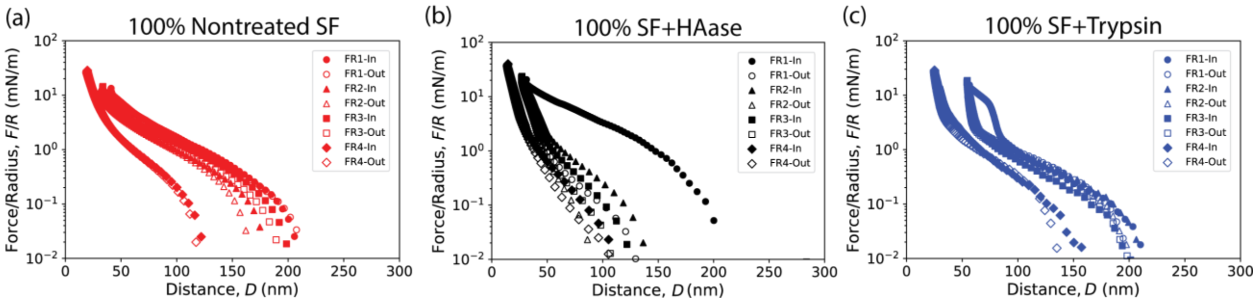


**SI Figure 4. Normal interaction forces between two mica surfaces across nondiluted SF.** All conditions were purely repulsive, and no adhesion was measured. (a) Nontreated SF showed little hysteresis between loading (in) and unloading (out), as well as between compression cycles (FR1-3). (b) SF+HAase showed significant hysteresis between, and (c) SF+Trypsin**.**

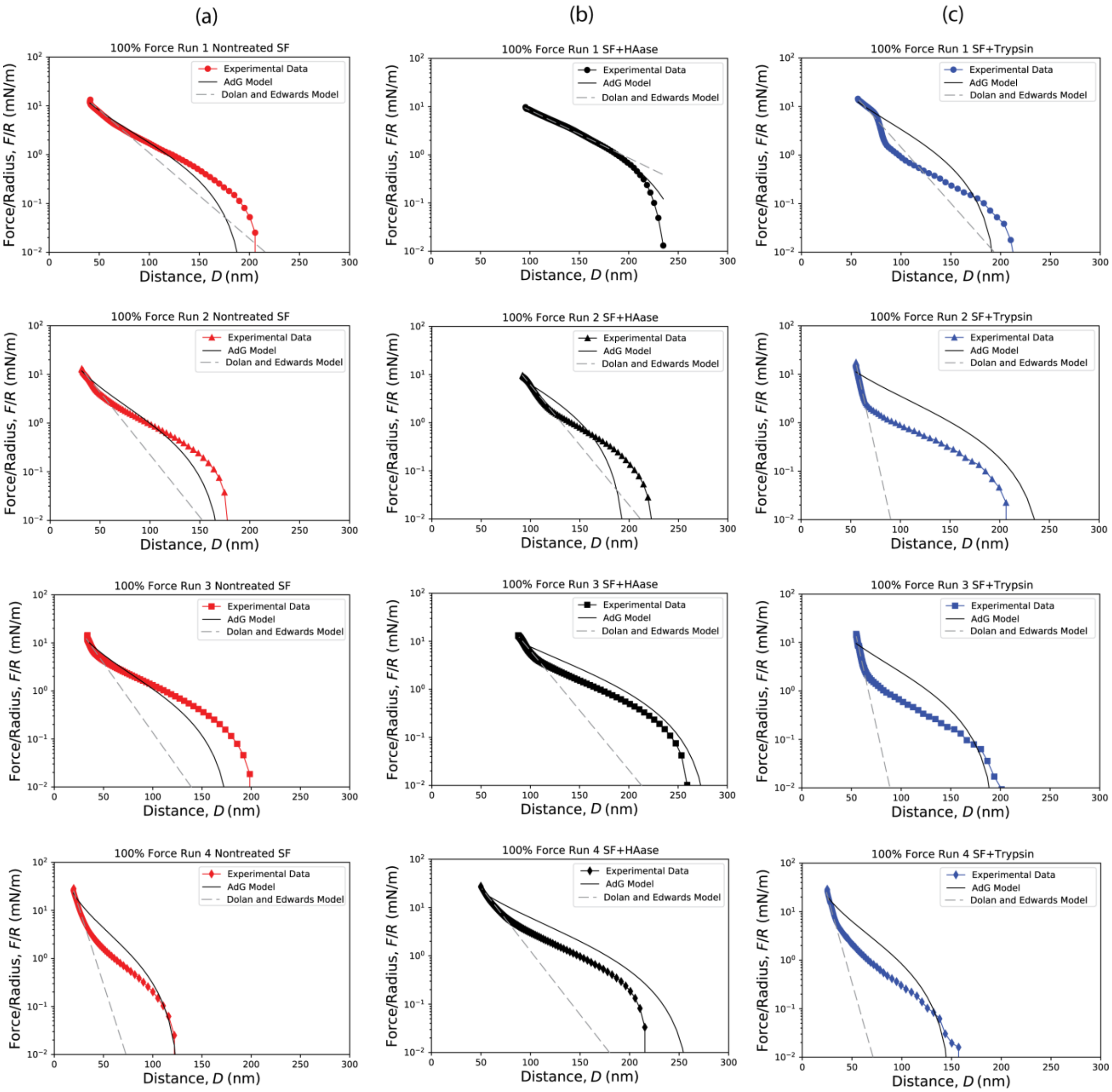


**SI Figure 5. Experiment, Alexander-de Gennes (AdG), and Dolan and Edwards (DnE) model comparison for 100% nontreated SF, FS+HAase, and SF+Trypsin.** Comparison between experimental normal interaction forces and best fi values for the AdG theory of end-grafted polymer brushes (eq. 5) and DnE model for two mica surfaces across SF. Note that the experimental data, AdG, and DnE models are shown in semilogarithmic plots.

**SI Table 4. Parameters obteined from non-linear least-square fits for AdG and DnE models for 100% SF films.** Effective brush thickness (*L*) and grafting distance (*s*) from the AdG brush model (eq. 5), and radius of gyration ($R_g$) and pre-factor (*A*) from the DnE model (eq. 6) used to fit the interaction forces between two mica surfaces across SFs for Force Runs 1-3 in.

| Condition | Alexander-de Gennes model, eq. 5 | | | Dolan and Edwards model, eq. 6 | | |
|---|---|---|---|---|---|---|
| | Effective brush thickness, *L* (nm) | Effective grafting distance, *s* (nm) | Fit $r^2$ value | Effective coil size, $R_g$ (nm) | Prefactor, *A* (mN/m) | Fit $r^2$ value |
| **Nontreated SF** | 93 ± 6 | 0.59 ± 0.01 | 0.98 ± 0.01 | 19 ± 5.0 | 84 ± 26 | 0.98 ± 0.01 |
| **SF + HAase** | 125 ± 23 | 0.47 ± 0.08 | 0.95 ± 0.02 | 663 ± 1100 | 1320 ± 1070 | 0.99 ± 0.01 |
| **SF + Trypsin** | 107 ± 14 | 0.54 ± 0.07 | 0.88 ± 0.07 | 9.5 ± 8.0 | 1,228,045 ± 1,087,503 | 0.99 ± 0.01 |
| **Bovine SF*** | 60 ± 13 | 1.15 ± 0.23 | NA | NA | NA | NA |

*From reference [15]

### Normal interaction forces between mica surfaces across diluted (5%) SF

Next, based on the Langmuir adsorption isotherm results, we asked how partially formed films would respond to confinement. We investigated the interaction forces of mica across 5% nontreated SF, SF+HAase, and SF+Trypsin using the SFA. Figure 5 and SI Figure 6 show representative force-distance profiles. Like what observed for nondiluted (100%) films, the force-distance profiles were purely repulsive, and the range and the magnitude of the repulsion varied with treatment and compression cycle. Nontreated SF and SF+HAase force-distance profiles were alike qualitatively and quantitatively. The onset of repulsion, $D_0$, for both treatments was measured to be ~95 nm, reaching a HW at ~10 nm. $D_0$ shifted to lower values with each subsequent compression cycle, to ~75 nm, and ~50 nm. The HW value remained constant for both treatments after a second and third compression at ~10 nm. This was not the behavior shown by SF+Trypsin, where $D_0$ for the first compression cycle (Force Run 1, Figure 5(a)) was ~200 nm, reaching a HW at ~50 nm. $D_0$, and HW values decreased with each subsequent compression cycle. For $D_0$, values decreased to ~90 nm and ~60 nm, and for HW to ~25 nm and ~10 nm, for Force Run 2 and 3, respectively. While little to no hysteresis was observed for 5% nontreated SF and for 5% SF+HAase either between in (loading) and out (unloading) or between cycles (Force Run 1-3), 5% SF+Trypsin showed considerable hysteresis, between in/out (loading/unloading) and between cycles (Force Runs). Interestingly, $D_0$ for Force Run 2 and 3 were very close to the termination of interaction of Force Run 1 and 2, respectively. This observation suggests that the film has been plastically deformed.

Repeating the experimental procedures introduced for nondiluted SF (100%) and to test if molecular film re-arrangement continued with higher applied normal forces ($F/R$ > 200 mN/m), we manually compressed nontreated SF, SF+HAase, and SF+Trypsin films until visible surface deformation/flattening occurred, followed by a final motor-controlled loading-unloading cycle, 10 minutes after the manual compression. All conditions experienced plastic/viscoelastic deformation, observed by a shift in $D_0$ to smaller distances. In this configuration, interestingly, all three conditions responded similarly to the dynamic force-distance measurement after the manual compression. Nontreated SF, SF+HAase, and SF+Trypsin $D_0$ decreased to ~50-70 nm, while the HW remained at ~10 nm, Figure 5(d), with minimal hysteresis between in/out (loading/unloading) profiles.

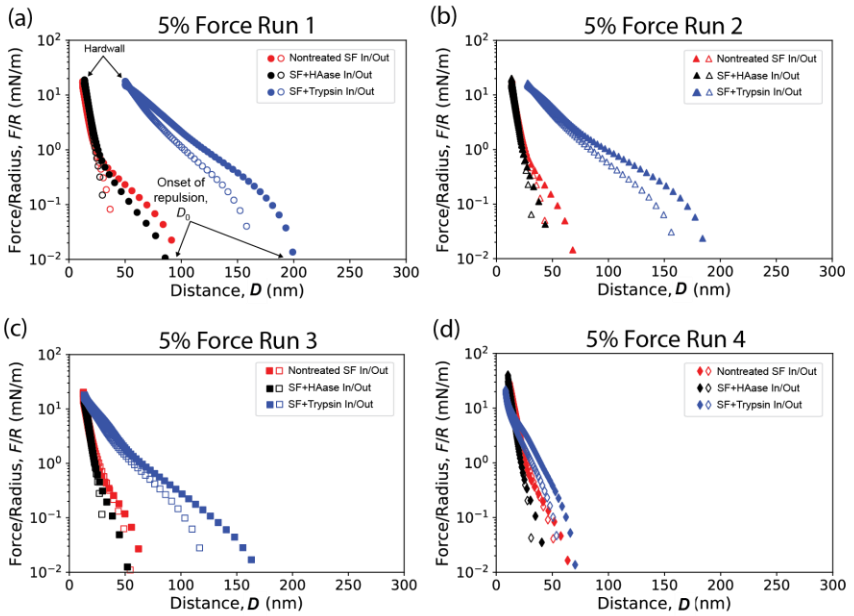


**Figure 5. Comparison of normal interaction forces between two mica surfaces across 5% diluted synovial fluid.** All conditions were purely repulsive, and no adhesion was measured. Profiles of the loading (in, filled symbols) and unloading (out, open symbols) for (a) first cycle, (b) second cycle, (c) third cycle, and (d) forth cycle of nontreated SF, SF+HAase, and SF+Trypsin**.**

We used the same two models, the AdG (eq. 5) and the DnE (eq.6) employed in the previous section to understand the repulsive behavior of the diluted films. Experimental data and fitted parameters obtained from non-linear least-squares are shown in SI Figure 7 and summarized in SI Table 4. Frist, we used the AdG model (eq.5) to obtain the average effective brush thickness (*L*) and average effective grafting distance (*s*) values. For nontreated SF, *L* and *s* were 49 ± 8 nm and 0.6 ± 0.1 nm, values that were similar in order of magnitude to SF+HAase, 33 ± 18 nm and 0.4 ± 0.2 nm for *L* and *s*, respectively. Interestingly, for SF+Trypsin, larger *L* and *s* values were observed, 107 ± 9 nm and 0.7 ± 0.3 nm, respectively. These values are very close to what measured for nondiluted SF+Trypsin (*L* = 107 ± 14 nm; *s* = 0.54 ± 0.07 nm).

Second, we used the Dolan and Edwards model (eq.6) to acquire the effective coil size, $R_{eff}$, and pre-factor *A* to model the SF films as end-grafted polymers in a mushroom configuration. For nontreated SF, $R_{eff}$ and *A* were 3.8 ± 0.2 nm and 462 ± 78 mN/m and for SF+HAase 3.6 ± 0.2 nm and 690 ± 78 mN/m, respectively. Not surprisingly, based on the previous AdG results, the $R_{eff}$ and *A* values for SF+Trypsin were 22.5 ± 4.6 nm and 63 ± 37 mN/m, an order of magnitude higher for $R_{eff}$ compared to nontreated SF and SF+HAase, and an order of magnitude lower for the pre-factor *A* compared to nontreted SF and SF+HAase. Based on the coefficient of determination ($r^2$) from our least-squares regression of the AdG and DnE models, summarized in SI Table 4, we observe that the prediction of the confinement force normalized by the radius of curvature as a function of the separation distance is more accurately described as an end-grafted polymer in the mushroom regime.

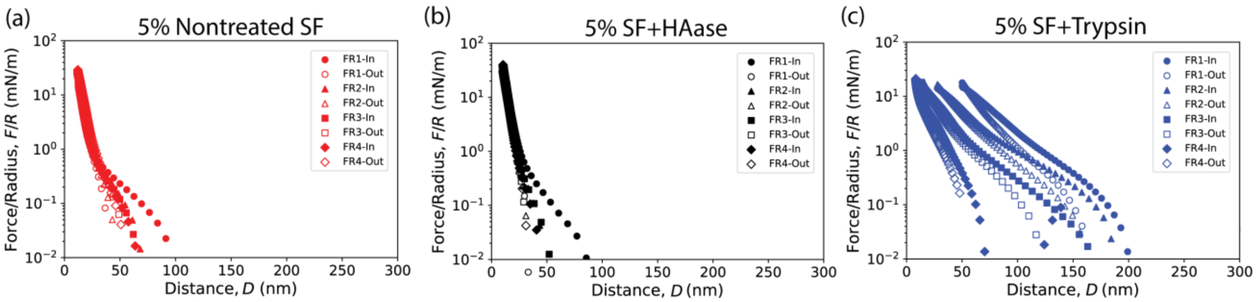


**SI Figure 6. Normal interaction forces between two mica surfaces across bovine synovial fluid.** All conditions were purely repulsive, and no adhesion was measured. (a) Nontreated SF showed little hysteresis between loading (in) and unloading (out), as well as between compression cycles (FR1-3). (b) SF+HAase showed significant hysteresis between, and (c) SF+Trypsin**.**

**SI Table 5. Parameters obtained from non-linear least-square fits for AdG and DnE models for 5% SF films.** Effective brush thickness (*L*) and grafting distance (*s*) from the AdG brush model (eq. 5), and radius of gyration ($R_g$) and prefactor (*A*) from the DnE model (eq. 6) used to fit the interaction forces between two mica surfaces across SFs for Force Runs 1-3 in.

| **Condition** | **Alexander-de Gennes model, eq. 5** | | | **Dolan and Edwards model, eq. 6** | | |
|---|---|---|---|---|---|---|
| | **Effective brush thickness, *L* (nm)** | **Effective grafting distance, *s* (nm)** | **Fit $r^2$ value** | **Effective coil size, $R_g$ (nm)** | **Pre-factor, *A* (mN/m)** | **Fit $r^2$ value** |
| **Nontreated SF** | 49.0 ± 8.0 | 0.6 ± 0.1 | 0.93 ± 0.005 | 3.8 ± 0.2 | 462 ± 78 | 0.99 ± 0.001 |
| **SF + HAase** | 33.0 ± 18.0 | 0.4 ± 0.2 | 0.93 ± 0.005 | 3.6 ± 0.2 | 690 ± 78 | 0.99 ± 0.001 |
| **SF + Trypsin** | 107.0 ± 9.0 | 0.7 ± 0.3 | 0.99 ± 0.001 | 22.5 ± 37 | 4.6 ± 37 | 0.99 ± 0.001 |

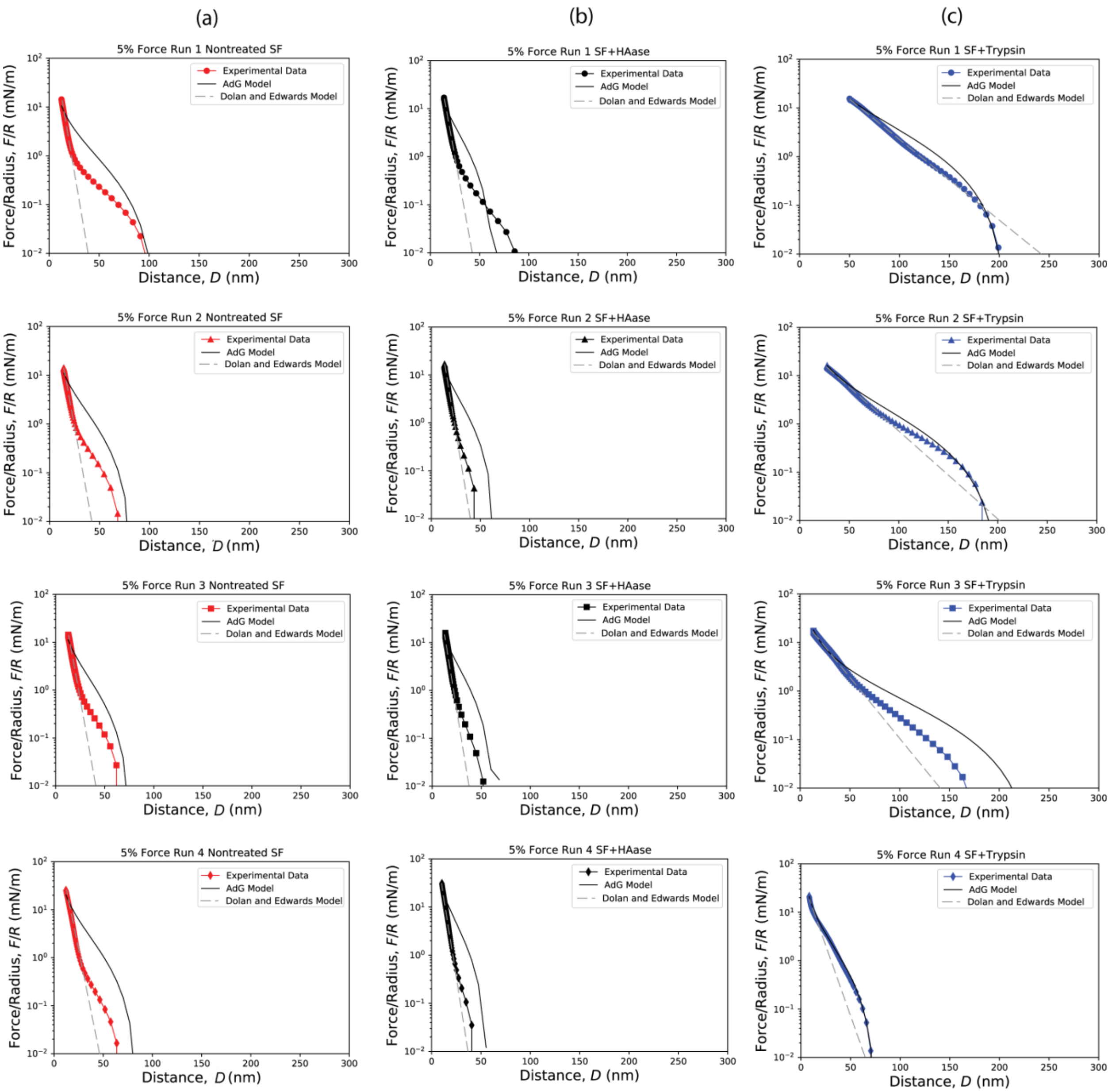


**SI Figure 7. Experiment, Alexander-de Gennes (AdG), and Dolan and Edwards (DnE) model comparison for 5% nontreated SF, FS+HAase, and SF+Trypsin.** Comparison between experimental normal interaction forces and best fi values for the AdG theory of end-grafted polymer brushes (eq. 5) and DnE model for two mica surfaces across SF. Note that the experimental data, AdG, and DnE models are shown in semilogarithmic plots.

**SI Table 6. Parameters obtained from non-linear least-square fits for AdG and DnE models for 5% SF films.** Effective brush thickness (L) and grafting distance (s) from the AdG brush model (eq. 5), and radius of gyration ($R_g$) and prefactor (A) from the DnE model (eq. 6) used to fit the interaction forces between two mica surfaces across SFs for Force Runs 1-3 in.

| **Condition** | **Alexander-de Gennes model, eq. 5** | | | **Dolan and Edwards model, eq. 6** | | |
|---|---|---|---|---|---|---|
| | **Effective brush thickness, *L* (nm)** | **Effective grafting distance, *s* (nm)** | **Fit $r^2$ value** | **Effective coil size, $R_g$ (nm)** | **Pre-factor, *A* (mN/m)** | **Fit $r^2$ value** |
| **Nontreated SF** | 49.0 ± 8.0 | 0.6 ± 0.1 | 0.93 ± 0.005 | 3.8 ± 0.2 | 462 ± 78 | 0.99 ± 0.001 |
| **SF + HAase** | 33.0 ± 18.0 | 0.4 ± 0.2 | 0.93 ± 0.005 | 3.6 ± 0.2 | 690 ± 78 | 0.99 ± 0.001 |
| **SF + Trypsin** | 107.0 ± 9.0 | 0.7 ± 0.3 | 0.99 ± 0.001 | 22.5 ± 37 | 4.6 ± 37 | 0.99 ± 0.001 |

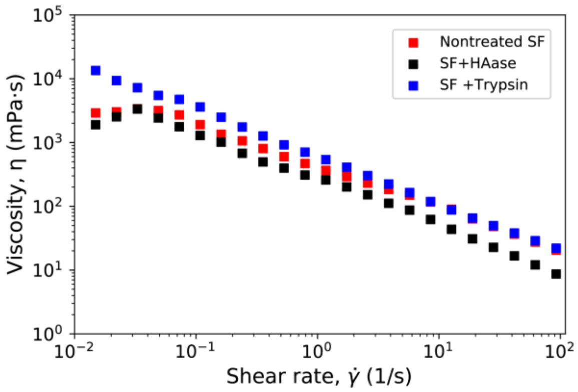


**SI Figure 8. Bulk viscosity of SFs.** Nontreated SF, SF+HAase, and SF+Trypsin as a function of shear rate.

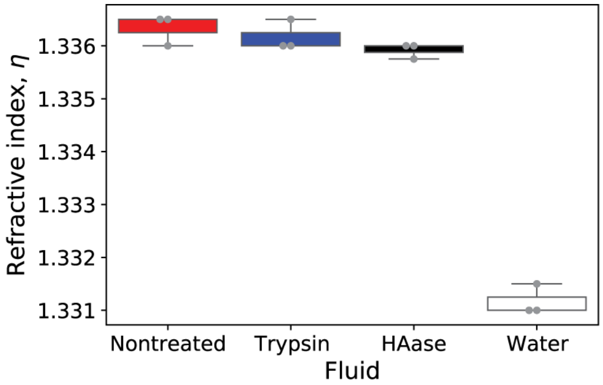


**SI Figure 9. Bulk refractive index of SF.** Digestion of HA and proteins did not change the refractive index of SF. Water refractive index value is shown as a control.
